# Measuring Neuronal Signals with Microelectrode Arrays: A Finite Element Analysis

**DOI:** 10.1101/2020.06.07.139014

**Authors:** R Bestel, U van Rienen, C Thielemann, R Appali

## Abstract

**Objective:** Measuring neuronal cell activity using microelectrode arrays reveals a great variety of derived signal shapes within extracellular recordings. However, possible mechanisms responsible for this variety have not yet been entirely determined, which might hamper any subsequent analysis of the recorded neuronal data. For an investigation of this issue, we propose a computational model based on the finite element method describing the electrical coupling between an electrically active neuron and an extracellular recording electrode in detail. This allows for a systematic study of possible parameters that may play an essential role in defining or altering the shape of the measured electrode potential. Our results indicate that neuronal geometry and neurite structure, as well as the actual pathways of input potentials that evoke action potential generation, have a significant impact on the shape of the resulting extracellular electrode recording and explain most of the known signal shape variety.

## I. Introduction

**A**nalyzing the electrical activity of neuronal cultures offers the opportunity to investigate communication in neuronal networks as well as the influence of various external stimuli. Compared to experiments *in vivo, in vitro* studies based on microelectrode arrays (MEA) often allow for more defined experimental setups. Electrical activity, i.e., neuronal action potentials (AP), can be either derived with standard MEA or using more recently developed high-density MEA that offer better spatial resolution [1], [2]. This data can be analyzed, e.g., in terms of global network activity, synchrony, or functional connectivity. Such methods rely on accurate detection of neuronal AP, also called spikes in their derived form that are present in a recorded signal. In the case of multiple neurons contributing to an electrode recording, each spike should be traced back to the cell that generated the respective AP to avoid errors in the subsequent data analysis. In this context, a complex issue arises, as measured spikes show significant shape variations [3], [4], while the origin of this phenomenon, as well as the relevant factors behind it, are still not entirely understood.

One possible approach to address this issue is via computational *in silico* models. Based on earlier neuron models [5], [6], [7], [8], we have conducted a simulation study to evaluate the effect of neuron morphology on the resulting electric potentials in extracellular space during neuronal AP-generation and -propagation [12].

In comparable works, e.g. [9], [10], [11], a cable equation-based approach was used, and the electrode signals were calculated using point- and line-source approximation. Yet, while cable equation-based models allow for the simulation of larger neuronal structures and multiple neurons in small networks, its applicability is fundamentally limited to rotational symmetric structures. In consequence, the accuracy of this approach decreases compared to the EQS-approach for small, irregular geometries, as, e.g., neurons that are adherent on an extracellular surface or an MEA-electrode (see [12]). In addition, point and line-source approximation may not account for all possible effects of the setup in extracellular space, as possible interdependences may be omitted.

In our previous work (see [12]), we have shown that both geometry and ion channel distribution have a significant impact on the EAP that is created during neuronal AP-generation and subsequent -propagation. Furthermore, the simulations indicated that only minor geometric changes are necessary to affect the resulting extracellular potential distribution.

Since the geometry of the neuron determines the physical contact with an underlying surface, a suitable geometric approximation becomes crucial if the measurement of a neuronal AP using an extracellular MEA-electrode is to be reproduced within a mathematical model. For this purpose, the results of our previous work presented in [12] showed fundamental advantages of an electro-quasistatic-based (EQS) approach compared to a cable equation-based approach to describe neuronal AP-propagation, as it allows for irregular geometric shapes and therefore for a much more accurate approximation of neuronal geometry.

In an attempt to extend our previous simulation model, in this work, we introduce an extracellular electrode to the neuron model and further refine the geometric and electrophysiological representation of the neuron. The finite element method (FEM) is used to describe the neuronal activity of a 3D neuron in an *in vitro* environment. Neuronal AP-generation is approximated with the Hodgkin-Huxley model and subsequent AP-propagation calculated using an EQS-based approach. Using this model-setup, the influence of electrode position concerning neuron geometry is evaluated and the effects of several parameters defining the electrical coupling between neuron and electrode are analyzed. Additionally, the influence of more complex neurite structures, as well as different origins to evoke neuronal AP-generation, is investigated.

Our results show that the signal shape measured by an extracellular electrode is highly dependent on its position to the neuron geometry. In addition, parameters such as neuron-electrode distance or electrode coverage, e.g., by packing glial cells, are found to have a significant influence on the measured signal amplitude. Finally, we show that the actual path of the AP, as well as the origin of evocation that induced initial AP-generation, has an essential influence on the shape of the extracellular electrode recording.

## II. Model description

Regardless of the actual neuron geometry, three distinct mathematical aspects need to be tackled. One task is the calculation of the changing intracellular potential *φ*_*i*_ due to AP-generation and subsequent propagation (see Fig. 1). Similarly, changes in the extracellular potential *φ*_*e*_ caused by neuronal electric activity that needs to be described. Finally, the ionic current *i*_*m*_ that is generated across the membrane during local AP-generation must be calculated.

**Fig. 1.**
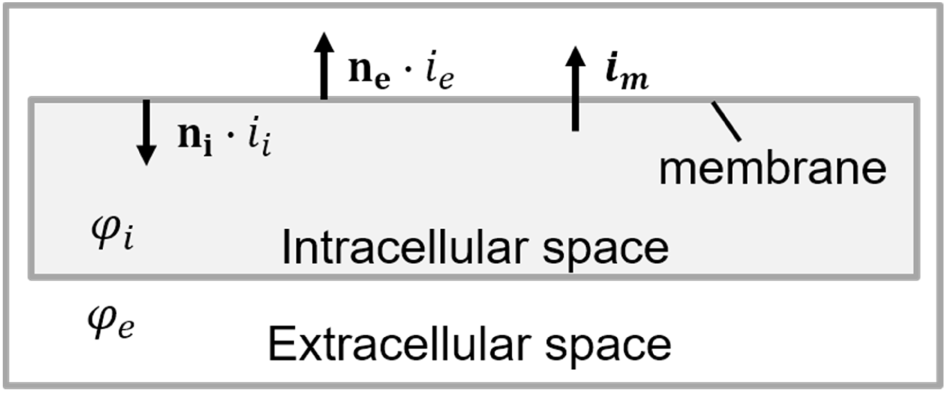
Schematic drawing of the mathematical parameters in the membrane model domain.

The transmembrane current *i*_*m*_ depends on the transmembrane potential *φ*_*m*_, defined as *φ*_*i*_ − *φ*_*e*_, and impacts the spatiotemporal evolution of the intra- and extracellular potentials *φ*_*i*_ and *φ*_*e*_ respectively. Consequently, a detailed description of the electrical activity, or in other words the AP-generation along the neuronal membrane is crucial for accurate simulation.

### A. Action potential generation with adapted Hodgkin-Huxley model

The Hodgkin-Huxley model is the most used mathematical approach to reproduce the AP-generation of neurons [13]. Its central equation describes the transmembrane current density *i*_*m*_ as the sum of ionic current densities *i*_*Na*_, *i*_*K*_ and *i*_*L*_ and the capacitive change of transmembrane potential *φ*_*m*_ as

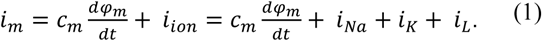

Transmembrane current densities due to *Na*^+^ and *K*^+^ ions are described by the terms *i*_*Na*_ and *i*_*K*_. The variable *i*_*L*_ combines a variety of secondary ions that only play a minor role. While the original model had been devised based on the electrophysiological properties of the giant squid axon, adapted models were proposed to capture the characteristics of mammalian neurons more accurately and take various temperature dependent effects into account [14], [15]. Similar to the original model each of the current densities is quantified in mA per m^2^ and generally calculated as

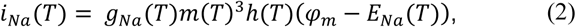

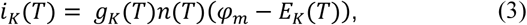

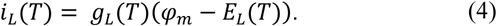

The parameters *E*_*Na*_, *E*_*K*_ and *E*_*L*_ describe the ion-channel-dependent reversal potentials and g_*Na*_, g_*K*_ and g_*L*_ the electrical membrane conductivities for the respective ion channel. The ionic currents *i*_*Na*_ and *i*_*K*_ are further governed by time- and space-dependent gating variables *m, h*, and *n*, describing the opening state of the corresponding ion channel. For the adapted model, all these parameters are dependent on the temperature *T* with the respective equations (A1-A3) shown in the appendix.

### B. EQS-approach

For describing AP-propagation and potential distribution in extracellular space, a system of EQS-based equations is used. To this end, a form of the continuity equation is implemented in both intra- and extracellular domains [6], [7], [12].

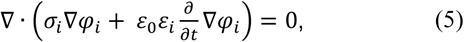

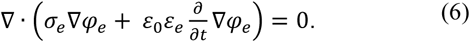

The neuronal membrane, which separates the two domains, is modeled using the thin-film approximation [16]. This means the membrane is not described as an additional volume domain but as two overlapping boundaries.

On each boundary, a position-dependent Neumann condition is implemented, which defines the transmembrane current density *i*_*m*_ along the respective normal vectors **n**_**i**_ and **n**_**e**_ into intra- and extracellular space

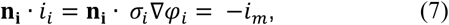

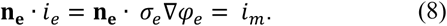

### C. Outer boundary conditions and electrode description

For the outer boundary at the bottom of the extracellular domain, a Neumann zero boundary condition is applied to model the insulating glass surface of an MEA with normal vector **n**_**s**_

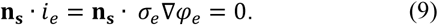

A circular area is defined on this surface representing the metal layer of an MEA electrode (see Fig. 2).

**Fig. 2.**
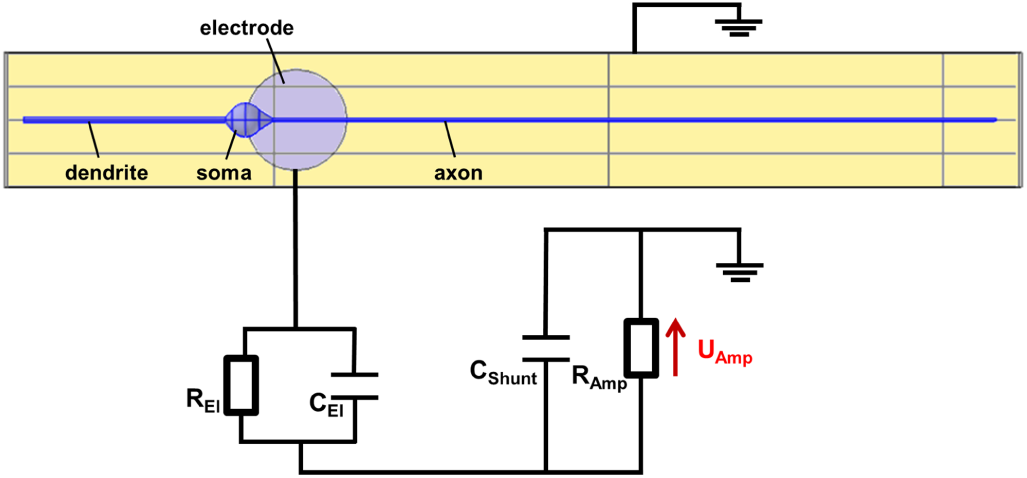
Schematic of the connection between the FEM model and the equivalent circuit description of the extracellular electrode (top view).

An electrical equivalent circuit based on previous mathematical models of [17], [18] describes the boundary condition of this electrode surface. The circuit capacitance *C*_*EL*_ and resistance *R*_*EL*_ describe the impedance of the electrode and the electrical double layer on top of the electrode surface. The capacitance *C*_*Shunt*_ accounts for losses across the MEA transmission lines. Finally, the resistance *R*_*Amp*_ defines the input resistance of the operational amplifier of the subsequent measuring system.

On the remaining outer boundaries, a Dirichlet zero condition is used to account for the reference electrode of the MEA, which sets the extracellular potential to zero.

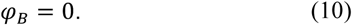

In order to create a complete electrical circuit, the respective zero potential condition is also defined for the description of the electrode circuit.

### D. Model geometry and electrophysiology

To evaluate the influence of neuronal geometry, three different neuron models I-III were created with increasing complexity. The general geometric parameters, such as soma diameter or axon and dendrite radius, were defined based on values presented in previous publications (see Table I).

**TABLE I.** Geometrical dimensions of Model I-III

| Quantity | Value |
| --- | --- |
| Soma diameter [19], [20] | 10 $\mu\text{m}$ |
| Axon diameter [21], [22] | 1 $\mu\text{m}$ |
| Axon hillock initial/end diameter | 3 $\mu\text{m}$ / 1 $\mu\text{m}$ |
| Dendrite diameter [19], [22] | 1.5 $\mu\text{m}$ |
| Dendrite tapering initial/end diameter | 1.5 $\mu\text{m}$ / 3 $\mu\text{m}$ |

General electrophysiological values of the neuron and electrical parameters of the extracellular medium were similarly taken from corresponding literature (see Table II).

**TABLE II.**
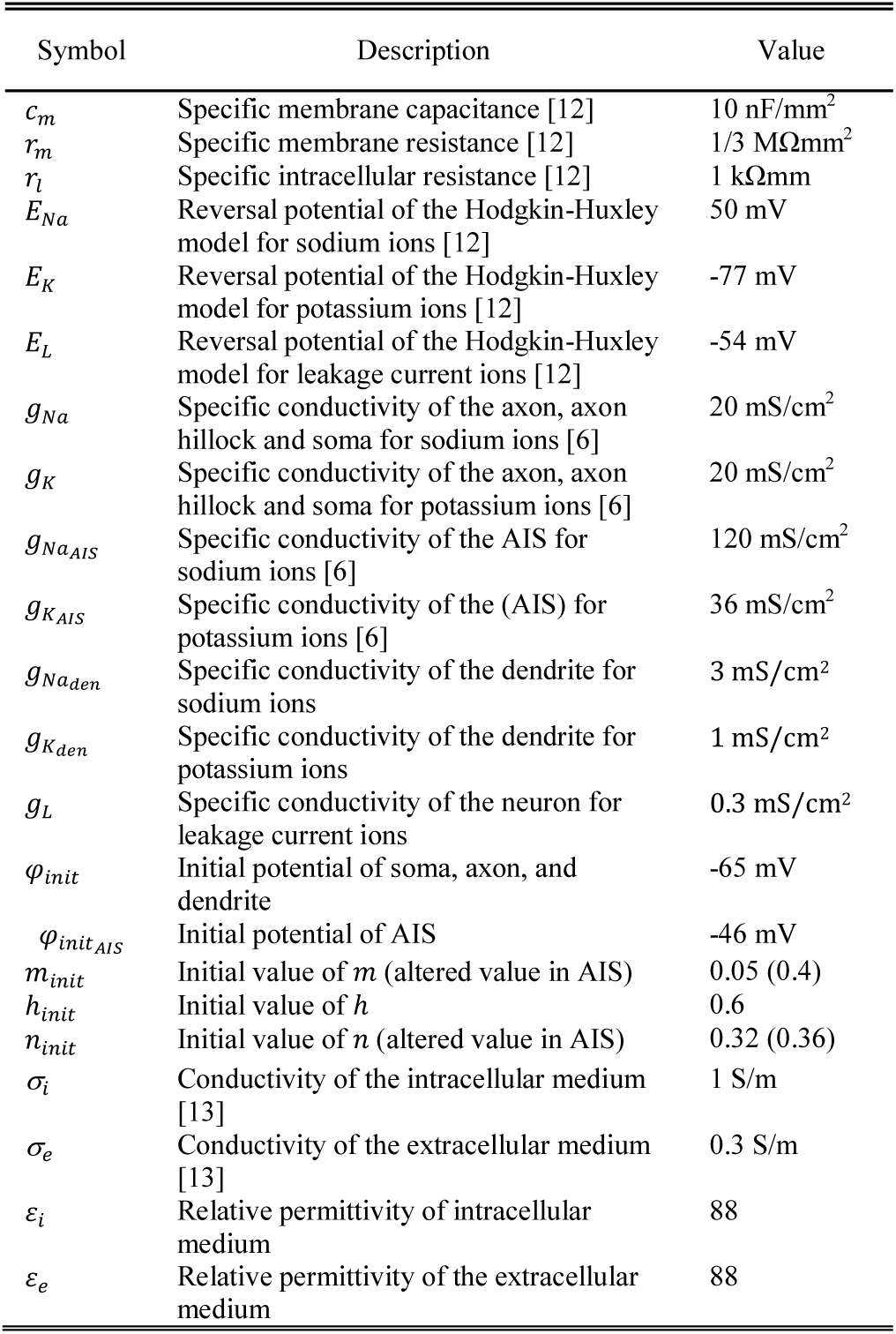
General electrophysiological parameters used in Model I-III

**Model I** consists of a simplified neuron geometry describing an adherent neuron on a MEA surface (see Fig. 3). The distance between the glass surface and the neuron is defined homogeneously as 100 nm [20]. The neuron model is generally comprised of geometric primitives, cylindrical for neurites, and spherical for the soma, while axon hillock and tapering from soma to dendrite are based on conical structures. These basic shapes are modified to approximate adherence of the cell on the glass substrate.

**Fig. 3.**
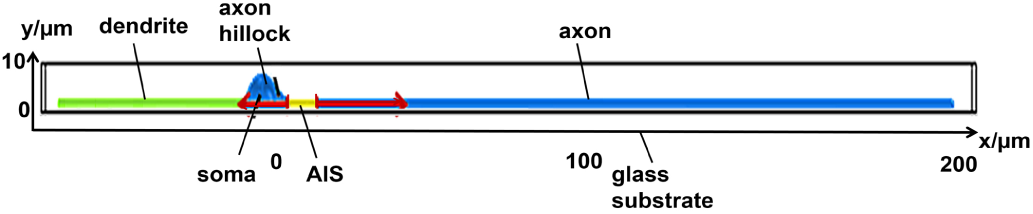
Schematic drawing of the model I (side view). The colors indicate different ion channel densities of the membrane (see Table II).

The surrounding extracellular space is defined as a cubic section of extracellular medium with electrical parameters of saline solution. Different densities of *Na*^+^ and *K*^+^ ion channels are imposed for each part of the neuron, e.g., soma, axon initial segment (AIS), and dendrite (see Table II). Initial AP-generation is induced at the AIS along the initial 10 µm of the axon. This is done by altering the initial values of the intracellular potential *φ*_*i*_ as well as the local ion channel gating parameters *m, n*, and *h*. Such a method has the advantage that possible artifacts caused by external current or voltage based stimulation can be avoided.

**Model II** features a refined neuron geometry with more realistic neurite patterns and multiple branches to approximate the shape of a bipolar neuron (see Fig. 4). However, the dimensions of soma, axon, and dendrites, including the conical connections are not altered compared to model I. Likewise, electrophysiological properties, such as ion channel distribution and electrical membrane parameters as well as starting values of the AIS for initial AP-generation remain unchanged.

**Fig. 4.**
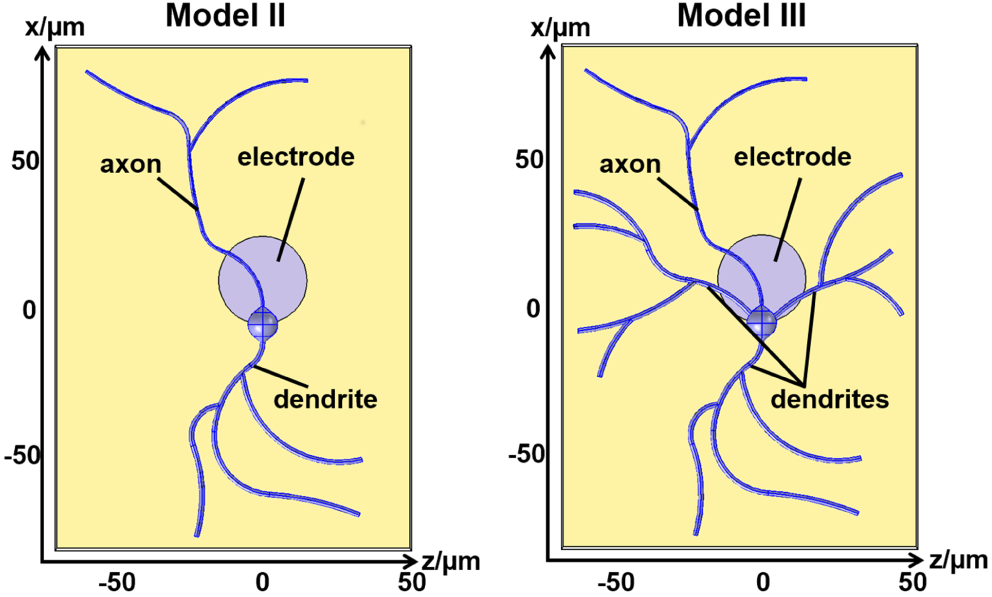
Schematic drawing of models II and III (top view).

For **Model III**, the geometry is further expanded by additional basal dendrites in order to reproduce the typical shape of a cortical pyramidal neuron (see Fig. 4). Similar to model II, all general geometrical and electrical properties are kept constant in order to accurately analyze the effect of neurite geometry on derived electrode signals.

### E. Computation

All models were created and simulated using the software COMSOL Multiphysics® 5.2 (COMSOL AB, Stockholm, Sweden). For spatial discretization, a tetrahedral mesh with second-order Lagrange elements was used.

The discretization of time-dependent quantities was done with an adaptive backward differentiation formula (BDF) scheme. Depending on the approximation error of the discrete solution in previous time steps, both the order of the BDF scheme and the size of the time step could be altered by the software for each iteration up to a maximum order of three and a maximum step of 10 µs. Nonlinearities in the resulting system were addressed using Newton’s method. Then, the linearized system was solved for each time step with the direct solver PARDISO using an allowed error tolerance of 1·10^−6^.

Simulations were carried out on a PC with two Intel® Xeon® E5-2687W v4 CPU with 24 cores, 256GB RAM, and a 64-bit operating system.

## III. Results

The initial analysis regarding the effect of parameters, such as neuron-electrode distance or electrode size on the resulting extracellular potentials and the respective electrode signals, is conducted on model I. Subsequently, models II and III are used to evaluate the general effect of neuron geometry onto derived electrode signals. Furthermore, the influence of different stimulus origins necessary to provoke AP-generation at the AIS is assessed based on the geometry of model III.

To ensure precise simulation, convergence concerning spatial and temporal discretization was determined for all models (see Table III).

**TABLE III.** Meshing and Computation Time of Model I-III

| Model | DOF<br>(with glial layer) | Computation time<br>(with glial layer) |
| --- | --- | --- |
| Model I | 403,000 (740,000) | 18 h (35 h) |
| Model II | 840,000 (-) | 31 h (-) |
| Model III | 1,740,000 (2,556,000) | 35 h (90 h) |

Because of their more complex geometries, the extended models I and III with additional glial layer required significantly higher degrees of freedom (DOF) for spatial discretization. For temporal discretization, a time step of 10 µs was found to be sufficient for all models. Considering computation time, the model I was solved within 18 h while results for the more complex Models II and III were calculated within 31 h and 35 h, respectively. Again, computation times of the extended models I and III with included glial layer coverage significantly increased to 35 h and 90 h. This is not only due to the increased spatial DOF but also due to the higher complexity of the resulting equation system, which is increased by the additional boundary conditions at the glial cell membranes.

To facilitate the model description, a general reference positon was defined for the MEA-electrode. It is positioned so that the center of the electrode has an offset of 15 µm with respect to the center of the soma in the x-direction (see Fig. 2). Any deviation from this reference positon is explicitely described for the respective simulation.

### A. Simplified neuron geometry (Model I)

The first step in analyzing the results of the model I is the evaluation of the resulting intra- and extracellular potentials during AP-generation and subsequent propagation. While the general properties and results of the individual modeling approach have been discussed extensively in our previous work [12], a brief overview, as well as the effects of the newly included extracellular electrode, are given in the following.

Based on the initial conditions, the AP is first generated in the AIS and then propagates along the axon (see Fig. 5). By using the modified Hodgkin-Huxley model of [15] at a temperature of 37° C, a propagation velocity of ∼100 mm/s is achieved. Furthermore, so-called back-propagation occurs with the AP being transmitted in the opposite direction into axon hillock, soma, and dendrite [24], [25], [26].

**Fig. 5.**
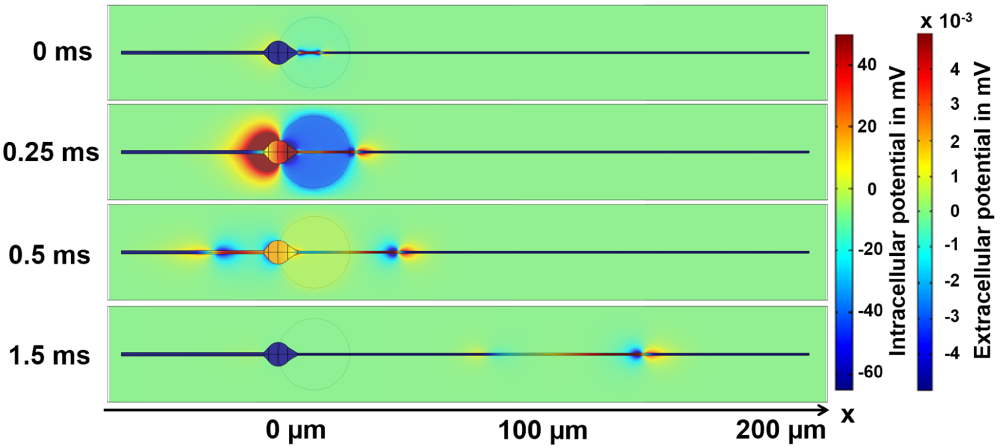
Intra- and extracellular potentials of model I. The electrode is positioned with an offset of 15 µm from the electrode center to the center of the soma (top view). Initiated in the AIS, the AP propagates along the axon in the x-direction as well as back into soma and dendrite. Shifts in the extracellular potential near the recording electrode during AP-propagation yield time-dependent changes of the electrode potential. Due to the high conductivity of the electrode, there is no significant potential gradient across the electrode surface at any given point in time.

In extracellular space, the most noticeable potentials are created during AP-propagation through the soma. The amplitude of these potentials is negative near axon hillock and AIS, whereas positive amplitudes are seen at the opposite side of the soma (see Fig. 5).

In short, this can be explained by the altered behavior of the ion channel kinetics at soma and axon hillock due to the change in diameter of intracellular space. This causes a more pronounced *Na*^+^ influx near the axon hillock and an increased *K*^+^ current near the junction of soma and apical dendrite (for details see [12]). In contrast, a uniformly shaped bipolar potential distribution is created in extracellular space during AP-propagation along axon and dendrite.

As can be seen best at 0.25 ms, an isopotential is created across the planar electrode surface for each point in time (see Fig. 6). The observed isopotential is caused by the high conductance of the metal electrode and shows the significant influence of the MEA-electrode on the extracellular potential in its vicinity. Due to the electrical properties of the extracellular medium and the boundary conditions in extracellular space, a time-varying potential distribution can be observed between the electrode and the reference potential at the outer boundaries of extracellular space.

**Fig. 6.**
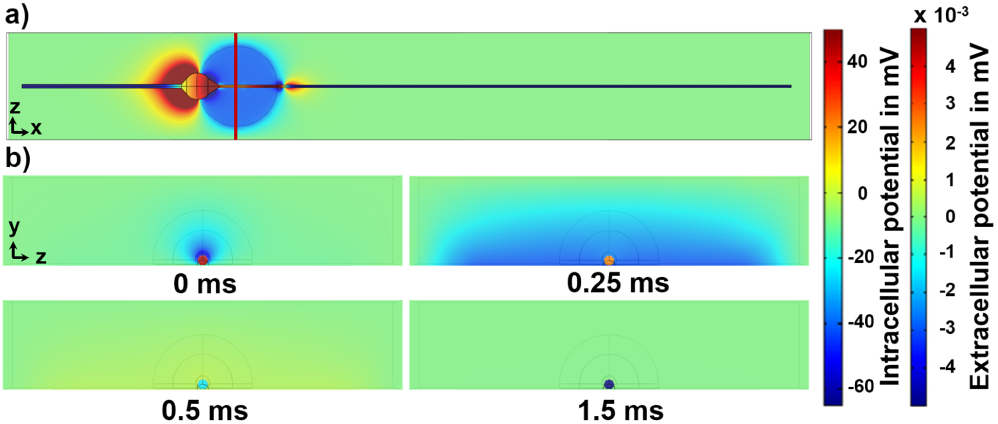
Time-dependent potential distribution near the extracellular electrode of the model I in front view with the electrode position indicated in a). b) A change in electrode potential caused by the electrical activity of the neuron yields a noticeable potential distribution in extracellular space, with values decreasing asymptotically towards the electrical ground, imposed at the top and side boundaries of the model.

Hence, a change in electrode potential, caused by the electrical activity of the adherent neuron, affects not only the entire electrode surface but also the potential distribution of the surrounding area in extracellular space.

Based on the inhomogeneous evolution of the extracellular potential seen in Figure 6, the derived voltage signals vary significantly with electrode position (see Fig. 7).

**Fig. 7.**
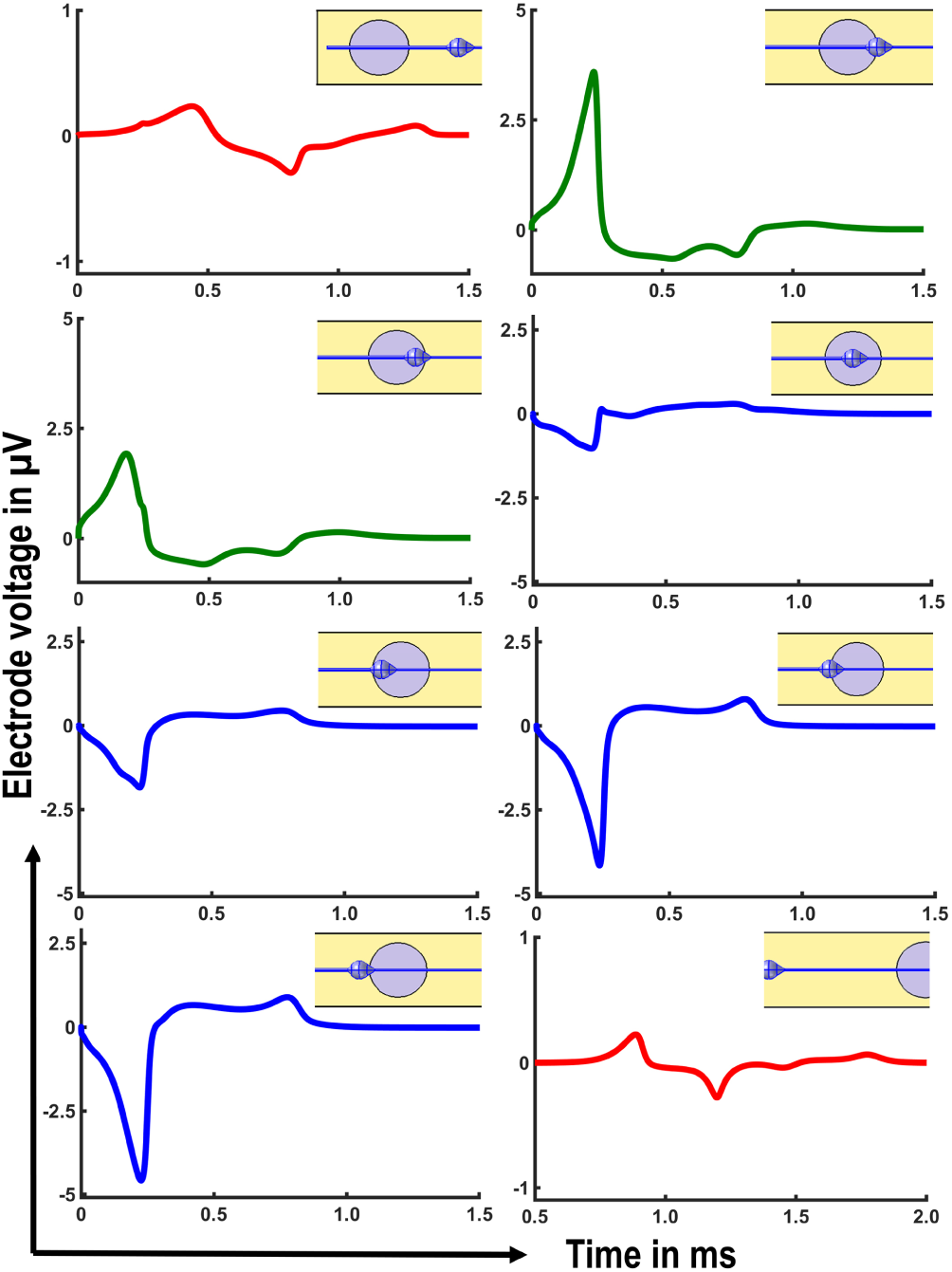
Electrode voltage at different recording sites along the neuron geometry of model I. The line colors indicate a different scale of the y-axis to improve visibility of each signal shape. Traces along axon and dendrite show a bipolar shape with amplitudes below ±1 µV. In contrast, signals recorded near the soma have noticeably higher amplitudes with values close to ±5 µV. Whereas traces recorded between soma and dendrite show a predominant positive amplitude, this is reversed on the opposite side of the soma near AIS and axon.

To highlight various shapes recorded for different electrode positions, y-axis scales are adapted for the shape types, which is emphasized by different line colors. In correspondence with respective extracellular potentials, electrode positions along axon and dendrite (red) yield small bipolar electrode signals. Near the soma, however, derived signals show noticeably different shapes with predominant negative amplitudes near AIS and axon hillock (blue) and positive amplitudes near the junction of soma and dendrite (green). With values of ± 5 µV signal amplitudes around the soma are noticeably higher than amplitudes measured along the neurites, which do not exceed ± 1 µV.

However, all derived signals are generally about one order of magnitude smaller than measured voltages with typical values around ± 30 - 100 µV (see [3], [18]). Compared to the geometry of the model I, which is only comprised of a single neuron to reduce computational complexity, *in vitro* cultures consist of a whole network of neurons often surrounded by a layer of supporting cells. Based on the results shown in Figure 6, one can assume that an additional cell layer significantly alters the potential distribution generated between the electrode surface and the reference potential at the outer boundaries of extracellular space. Such additional coverage changes the electric resistance between the electrode surface and the extracellular reference potential at the outer boundaries, which consequently leads to a higher electric potential at the electrode. Hence, the amplitude of the derived electrode signal is increased.

### B. Effect of insulating glial-layer on signal amplitude

For verification of a possible influence of electrode coverage on the amplitude of extracellular recordings, the model I is altered so that a layer of glial cells additionally covers the surface of the MEA-electrode (see Fig. 8).

**Fig. 8.**
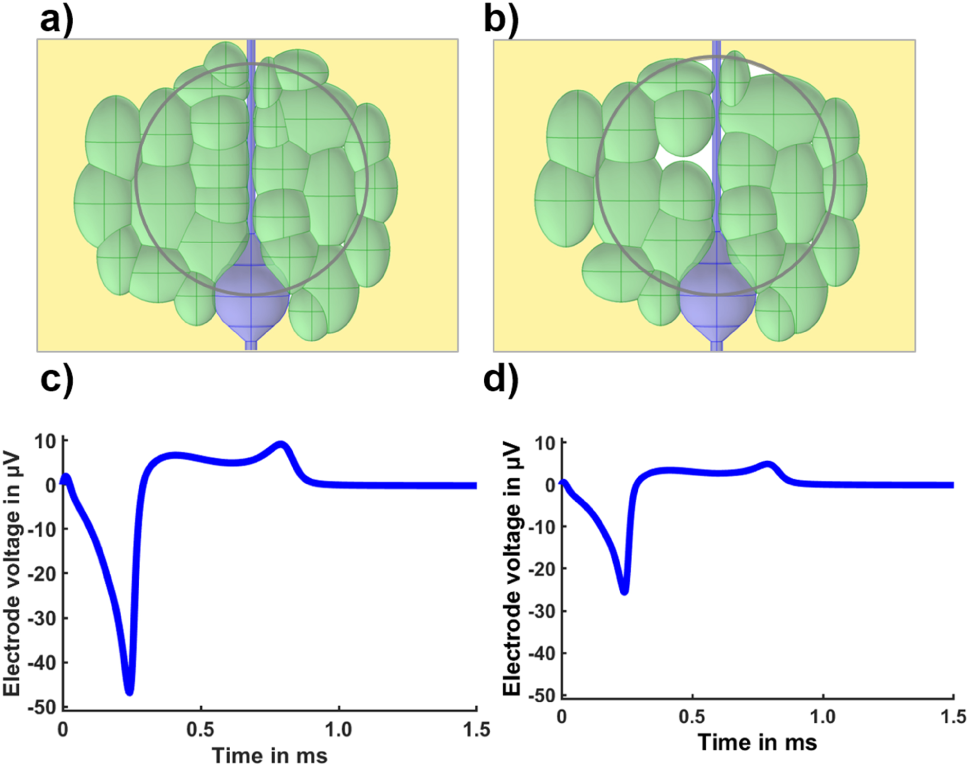
Effect of a glial layer on the amplitude of derived electrode signals. The glial layer (green) in a) fully covers the electrode surface (outlined in grey), whereas the layer that is shown in b) features some small voids (white areas). c) and d) While a full glial coverage yields an increase of the derived signal amplitude by approximately one order of magnitude, the effect is noticeably mitigated even by few voids inside the layer. The signal shape is not affected in either case.

Such cells are generally not considered to have an electrical activity similar to neurons [27], yet fulfill a supporting role in neural tissue and are also present in many *in vitro* cultures [28]. The introduced glial-layer is comprised of several ellipsoid-shaped cells with semi-axes ranging between 2-7 µm. Each glial cell is modeled with a resting potential of -90 mV, a passive cell membrane with a specific capacity 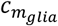 of 0.75 μF/*mm* and a specific leakage conductivity 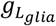 of 1· 10^−5^ S/*m* [29]. An electrode position with an offset of 15 µm (to the center of the soma in the *x-*direction) is chosen as reference. In agreement with previous results, a dense glial layer dramatically increases the resulting electrode signal amplitude by almost one order of magnitude to -45 µV (see Fig. 8a and 8c).

The 100 nm small gap that is formed by the electrode surface and glial layer in *y*-direction significantly increase the electric resistance between the electrode surface and extracellular mass potential. In consequence, the amount of electric current transmitted to the input impedance of the amplifier stage during the electrical activity of the adherent neuron is increased, while the amount of leakage current to the outer boundaries of extracellular space decreases, respectively. Hence, a noticeably higher electrode signal amplitude is achieved in case of additional electrode coverage. In addition, it can be shown that even small voids inside the layer cause a significant drop in amplitude (see Fig. 8b and 8d). However, the shape of the signal trace remains unaffected in both cases. Overall, the results of the modified model I verify that electrode coverage is a highly important factor. While non-covered areas of the electrode surface yield noticeable leakage currents, which significantly influence the amplitude of the derived electrode signal, reasonable signal amplitudes that are comparable with measured *in vitro* data can be achieved when imposing a dense coverage of the electrode with an additional glial layer.

### C. Effect of neuron-electrode distance and electrode size

As the simulations based on the model I have shown, both the shape and amplitude of the derived electrode signal are highly dependent on electrode position and neuron geometry (see Fig. 7). The impact of a possible offset in the *z-*direction, an increase in neuron-electrode distance in the *y-*direction, or the size of the MEA-electrode has not yet been investigated for the computational model. In the following, these three factors are addressed in a parameter study, and three particular situations of neuron-electrode coupling are evaluated:

Electrode setup 1: A planar MEA-electrode without additional electrode coverage

Electrode setup 2: A planar MEA-electrode with additional electrode coverage by a dense glial-layer (see Fig. 8a)

Electrode setup 3: A recessed MEA-electrode with a passivation layer of 1 µm thickness, which creates a respective cavity at the electrode site. No additional electrode coverage is imposed.

The previously used reference position of the electrode is retained with an offset of 15 µm in the *x-*direction to the center of the soma and a distance of 100 nm between the cell membrane and the electrode surface. Based on this reference position, the recessed electrode in electrode setup 3 yields a signal amplitude of -5.02 µV compared to -4.58 µV for electrode setup 1 and -45 µV for electrode setup 2 with additional glial layer coverage. The recessed electrode produces slightly higher signal amplitudes due to the small increase in electric resistance between the electrode surface and the reference potential at the boundaries of extracellular space compared to electrode setup 1. In the following parameter study, these voltages serve as a reference for each electrode setup, and the relative change of signal amplitude is evaluated (see Fig. 9).

**Fig. 9.**
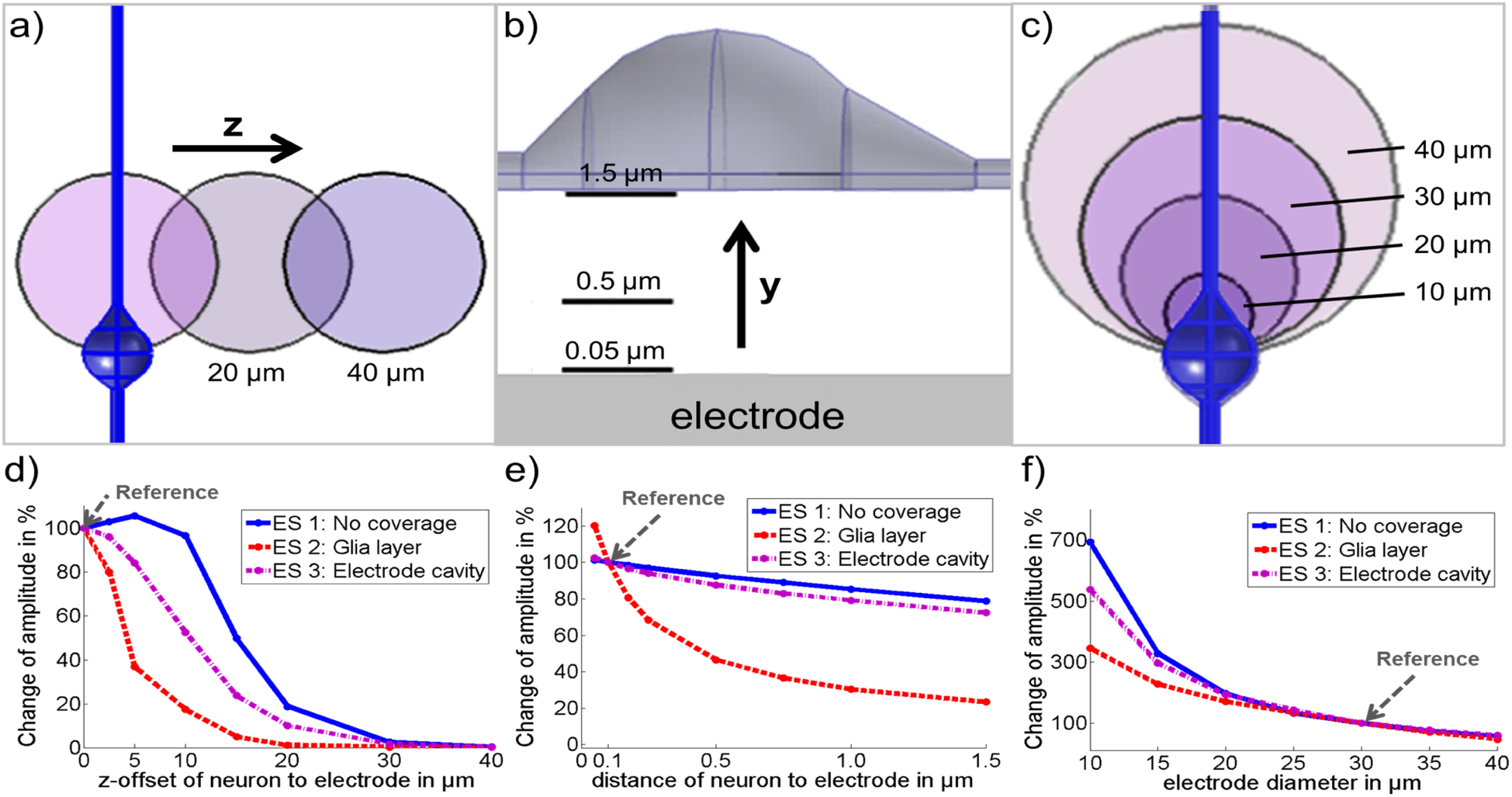
Evaluation of the influence of electrode displacement (a), neuron-electrode distance (b) and electrode size on the derived signal amplitude. d) In all of the three electrode setups (ES), the signal amplitude decreases dramatically with electrode displacement. The minor amplitude increase for a displacement of 5 µm in setup 1 can be explained by the extracellular potential distribution near the soma. e) For electrode setup 2, the derived signal amplitude decreases asymptotically with a more considerable distance of the neuron to the electrode. Due to the generally smaller signal amplitudes, the effect is noticeably weaker in case of setups 1 and 3, respectively. f) Signal amplitudes generally decrease with larger electrode diameter in all cases. Due to better conduction to the electrical ground in extracellular space, this effect is most pronounced for electrode setup 1

The signal amplitudes generally decrease asymptotically for all setups with increasing electrode offset in the *z*-direction (see Fig. 9d). Similar to previous results, the signal shape remains to be generally unaffected. However, the amplitude decrease differs between the models, with electrode setup 2 showing the steepest decline. Signal amplitude decreases to less than 10% of the reference amplitude at an offset of 15 µm, as the effect of additional electrode coverage is quickly lost for higher offsets.

Electrode setup 3 displays a slower reduction in signal amplitude, decreasing to ∼10 % at 20 µm, and electrode setup 1 shows the slowest decline, reaching values below 10 % of the original amplitude only well after a displacement of 20 µm.

In addition, the curve of electrode setup 1 shows a slight increase in the signal amplitude of 5 % for smaller displacements around 5 µm. This effect is due to the circular shape of the electrode in combination with the potential distribution near the neuron’s soma. While the positive potential generated near the connection between dendrite and soma is still partially captured for the reference electrode position, this already changes with small displacement in the *z-*direction. In consequence, the potential average across the electrode surface yields slightly higher negative amplitudes. However, this behavior is only visible due to the overall small signal amplitude of electrode setup 1 and cannot be seen in the two alternative setups.

Overall, all electrode setups generally show the expected decrease in amplitude in the case of a positional electrode displacement in the *z-*direction, which corresponds to more considerable distances between cell and electrode. Nonetheless, it is most evident for electrode setup 2, further highlighting the influence of current leakage due to an uncovered electrode surface. Furthermore, the cavity of electrode setup 3 makes it less susceptible to minor electrode displacement, even though not providing significantly higher signal amplitudes in general.

Considering higher distances of the cell membrane to the electrode surface in the *y-*direction, e.g., due to insufficient cell adhesion, electrode setup 1 and 3 yield a nearly linear decline with increasing distance (see Fig. 9e). In contrast, additional glial-coverage in electrode setup 2 leads to an asymptotic decline. In agreement with the previous results, this indicates that in setups 1 and 3, most of the electric current generated during AP-generation gets lost due to leakage currents rather than being transmitted to the input resistance of the amplifier. In consequence, increasing the distance between neuron and electrode and hence decreasing the electric resistance between the electrode surface and the reference potential only has a marginal effect. Yet, it has to be noted that the small effect of the distance between neuron and electrode for setup 1 and 3 is mainly caused by the low amplitude of the electrode signal for the reference positions. With the additional glial-layer coverage in electrode setup 2 however, signal amplitudes not only decrease more realistically with distance but also increase more significantly by more than 20% if the neuron-electrode distance is decreased to 0.05 µm.

Investigating the effect of different electrode sizes on the derived electrode signal amplitudes yields generally similar results for all three tested setups (see Fig. 9f). With a diameter of 30 µm as a reference, signal amplitudes overall increase noticeably for smaller diameters, while decreasing with larger electrode sizes. This increase is most significant for setup 1, yielding an amplitude that is almost seven times higher for an electrode diameter of 10 µm. For electrode setup 3, the signal amplitude is about 540% of the reference value, whereas setup 2 only shows an increase to 350%.

The different behavior of the electrode setups can again be explained by the leakage current that occurs in the case of uncovered electrode surfaces. The ratio between the area of the electrode surface covered by the neuron itself and the non-covered area is reduced when the electrode size is decreased (see Fig. 9c). This yields higher resistance to the reference potential at the outer boundaries of extracellular space and, thus a lower leakage current. In consequence, the amplitude of the resulting electrode signal increases with smaller electrode diameters. Since the amplitude loss due to leakage is highest in the case of electrode setup 1, the effect of electrode size is most noticeable for this setup.

### D. Physiological neuron geometry (Models II and III)

After a general analysis of the neuron-electrode coupling by employing the model I, the neuron geometry is revised by introducing models II and III. Both models feature shapes that are physiologically realistic and have multiple dendrites and axon branches. Therefore, the more complex geometries of models II and III as well as a comparison with the simplified model I allow for a comprehensive evaluation of the effect of neuron shape on the derived electrode signal.

In a first step, AP-generation is initiated similar to the model I in the AIS of models II and III. Subsequent AP-propagation along the axon as well as backpropagation into soma and dendrites occurs essentially identical in both models (see Fig. 10). Similar electric potentials are observed in extracellular space despite the geometric differences. Overall, results are in good agreement with the model I (see Fig. 6). The AP spreads into each of the neurite branches at the branching points and is consequently regenerated for further propagation. This causes higher extracellular potentials near these points, as additional ionic currents are required to ensure AP-propagation in all subsequent pathways.

**Fig. 10.**
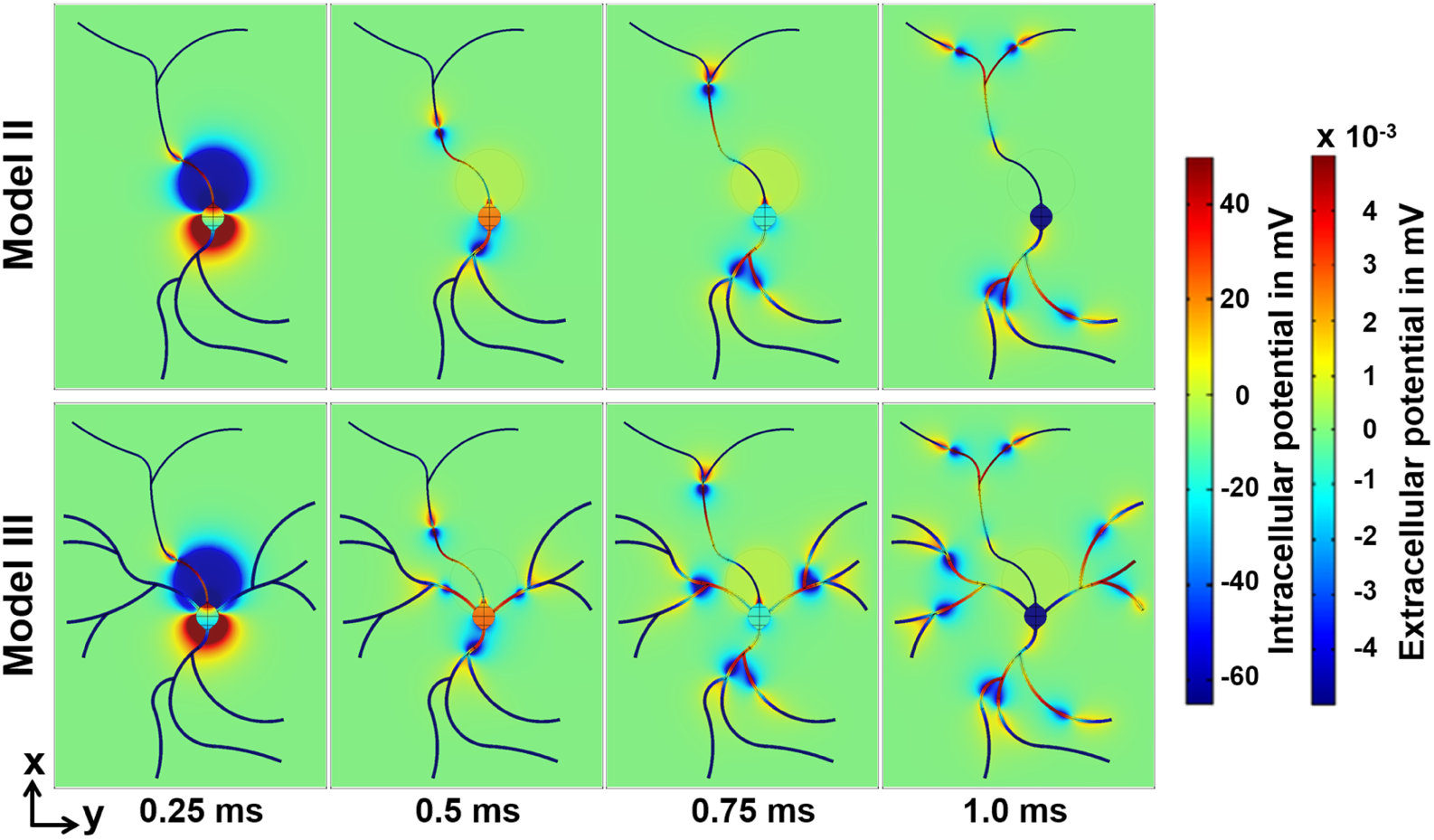
Intra- and extracellular potentials of models II and III (top view). Similar to the model I (see Fig. 5), the AP propagates in the *x*-direction along the axon as well as back into soma and dendrite for both models. AP-propagation occurs similarly in both models II and III, yet the additional dendritic structures of model III change the extracellular potential distribution in their vicinity.

Considering the derived electrode potential traces at various recording locations, models II and III show similar signals (see Fig. 11). Similar to figure 7, the y-axis is adapted for different signal shape types, which are hilghligted by varying line colors. Differences between the electrodes signal traces of the two models are found at positions close to the soma. These deviations are caused by the additional basal dendrites of model III that provide additional transmembrane currents and thus alter the extracellular potential in this area. Due to the same effect, electrode signals derived near branching points show increased amplitudes when compared to regular recording sites along axon and dendrite. In comparison with respective signals calculated using the model I, results are generally in good agreement in both shape and amplitude for respective electrode locations (see Fig. 5).

**Fig. 11.**
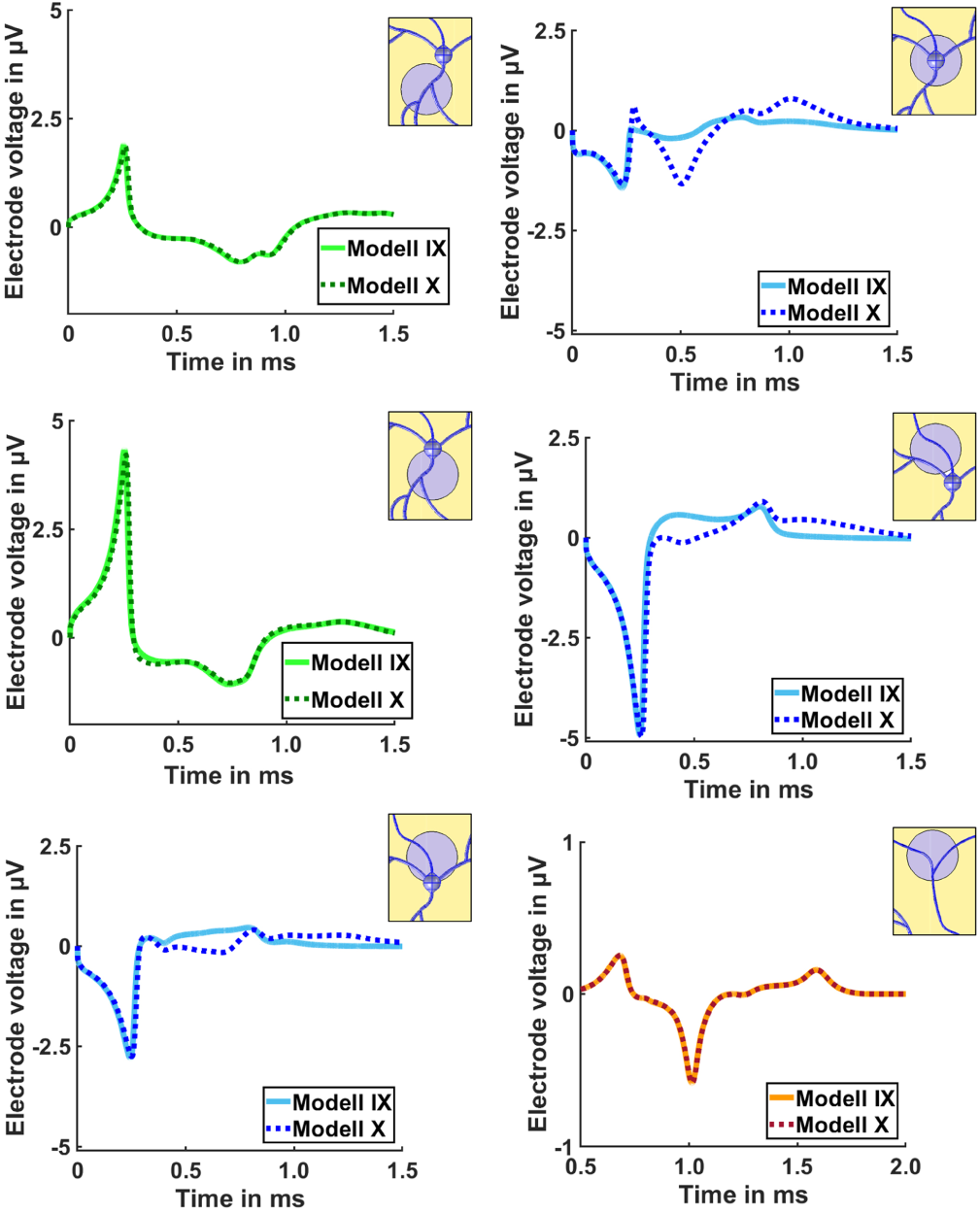
Electrode voltages derived at different recording sites on the neuron geometry of model II (solid line) and model III (dashed line). The line colors indicate a different scale of the y-axis to improve the visibility of each signal shape. While yielding mostly identical traces, the derived signal shapes differ near the soma, as the basal dendrites of model III generate additional ionic currents that alter the extracellular potential in this area.

In principle, all models yield electrode signals with negative amplitude near the axon hillock, whereas positive signal amplitudes are generated at the transition area of soma and apical dendrite. Along neurites, the extracellular electrode records a smaller, bipolar signal trace. Additional neurite branches like the basal dendrites of model III alter the derived signal shapes, as they provide additional ionic current sources and change the distribution of the local extracellular potential.

Considering the amplitudes of derived electrode signals, values for models II and III again do not exceed ±5 µV.

This can be explained Consequently, by imposing an additional glial layer, signal amplitudes are enhanced by about one order of magnitude (see Fig. 12).

**Fig. 12.**
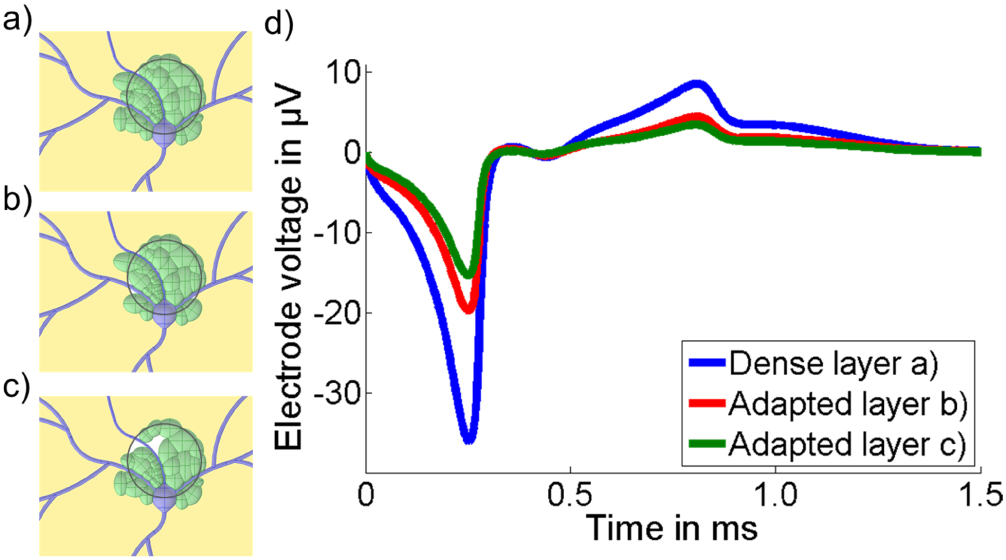
Derived electrode signals for model III with different glial layer setups, as shown in a) – c). d) In the case of dense electrode coverage, the signal amplitude is increased to -36 µV, which is more than five times higher compared to the original setup (see Figure 10). Similar to the model I even small voids inside the glial layer result in a significant decrease of signal amplitude. The signal shape, on the other hand, is not affected by either size or exact location of the voids.

Another aspect shown with the results of model III is that voids inside the glial layer affect mainly the amplitude of the recorded electrode signal but do not significantly change signal shapes. The exact location of the voids, e.g., near the basal dendrites in contrast to near axon hillock or soma, does not alter the shape of the signal.

### E. Origin of stimulating signal to evoke AP-generation

In all previous simulations, AP-generation and subsequent propagation were initialized in the AIS as spontaneous activity. Nonetheless, AP-generation is often evoked by incoming stimuli from surrounding neurons transmitted via the dendritic tree. Unlike the models I and II, model III offers different neurite pathways and input combinations to evoke AP-generation in the AIS. In order to analyze any effect of the origin of evocation on the resulting electrode signal, model III is simulated with four different input signal scenarios (see Fig. 13).

**Fig. 13.**
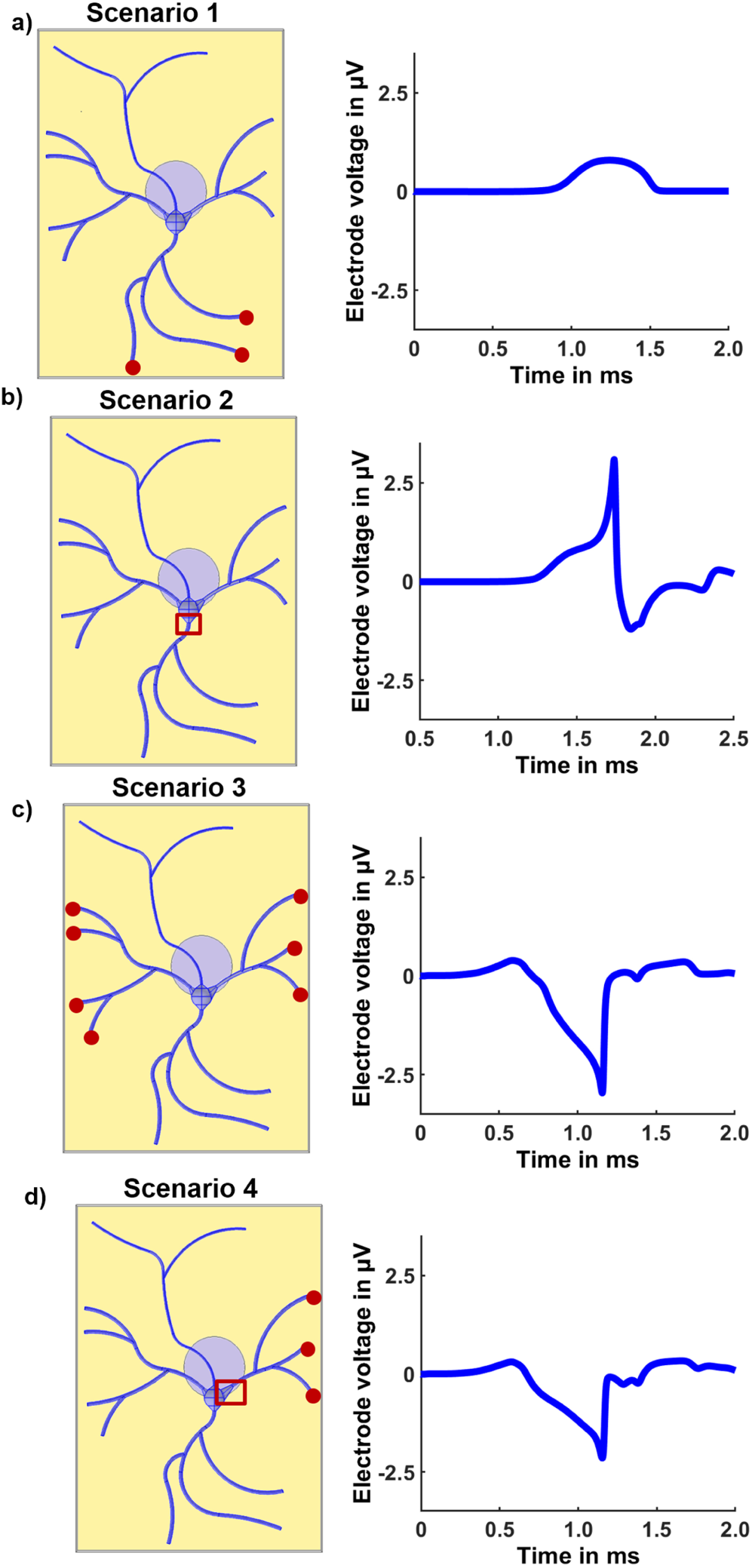
Resulting electrode signal for different stimulus origins concerning neuron geometry. a) – b) While the initiation of the AP generation in the AIS using the apical dendrite is not achieved for the original neuron geometry, expanding the junction between apical dendrite and soma facilitates stimulation. Based on the resulting potential distribution in extracellular space, this yields an electrode signal with a positive amplitude. c) – d) In contrast, stimulation via the basal dendrites results in a signal trace with negative amplitude. Similar to scenario 2, see b), an enlargement of the junction between dendrite and soma is necessary to initiate the AP generation with only a single basal dendrite.

AP propagation is initialized using a time-dependent potential trace as a boundary condition at the end of several dendrites. The potential trace is generated by the modified Hodgkin-Huxley model using the ion channel densities of the dendrite (see Table II) and mainly describes an incoming AP at the given boundary (see Fig. A1). For all scenarios, the reference electrode position with an offset of 15 µm in *x-* direction relative to the center of the soma is maintained.

In case of scenario 1, this boundary condition is solely imposed on the branches of the apical dendrite (see Fig. 13a). However, even though propagating along the dendrite simulation results show that the dendritic potential is unable to depolarize the larger volume of the soma and consequently AP-generation in the AIS fails.

The failure can be explained by the sudden increase of intracellular volume at the junction of dendrite and axon, which can cause failing AP-propagation (see [30]). In this context, the geometry at the junction between basal dendrite and soma is altered for scenario 2 to create a more gradual expansion of intracellular volume. After geometric alteration, the same dendritic input is sufficient to ensure the propagation of the AP through the soma, and hence AP-regeneration at the AIS is achieved (see Fig. 13b). In that, scenario 2 yields an electrode signal with an absolute positive amplitude of 2.5 µV.

Similar to scenario 1, the dendritic AP of a single basal branch of model III is insufficient to achieve AP-propagation across the soma. Nevertheless, with incoming evocation from both basal dendrite branches, AP-propagation, and subsequent regeneration can be achieved for scenario 3 (see Fig. 13c).

In contrast to scenario 2, this yields a monopolar electrode signal with an amplitude of -2.8 µV. In the scenario of scenario 4, the adaptation of the geometry at the junction of the basal dendrite facilitates AP-propagation into the soma, which allows for subsequent AP-propagation. The derived electrode signal is similar to scenario 3 but slightly reduced to -2.3 µV.

The explanation for the noticeable difference between electrode signals of scenario 2 compared to scenarios 3 and 4 can be found using the extracellular potential distribution due to AP-propagation (see Fig. 14).

**Fig. 14.**
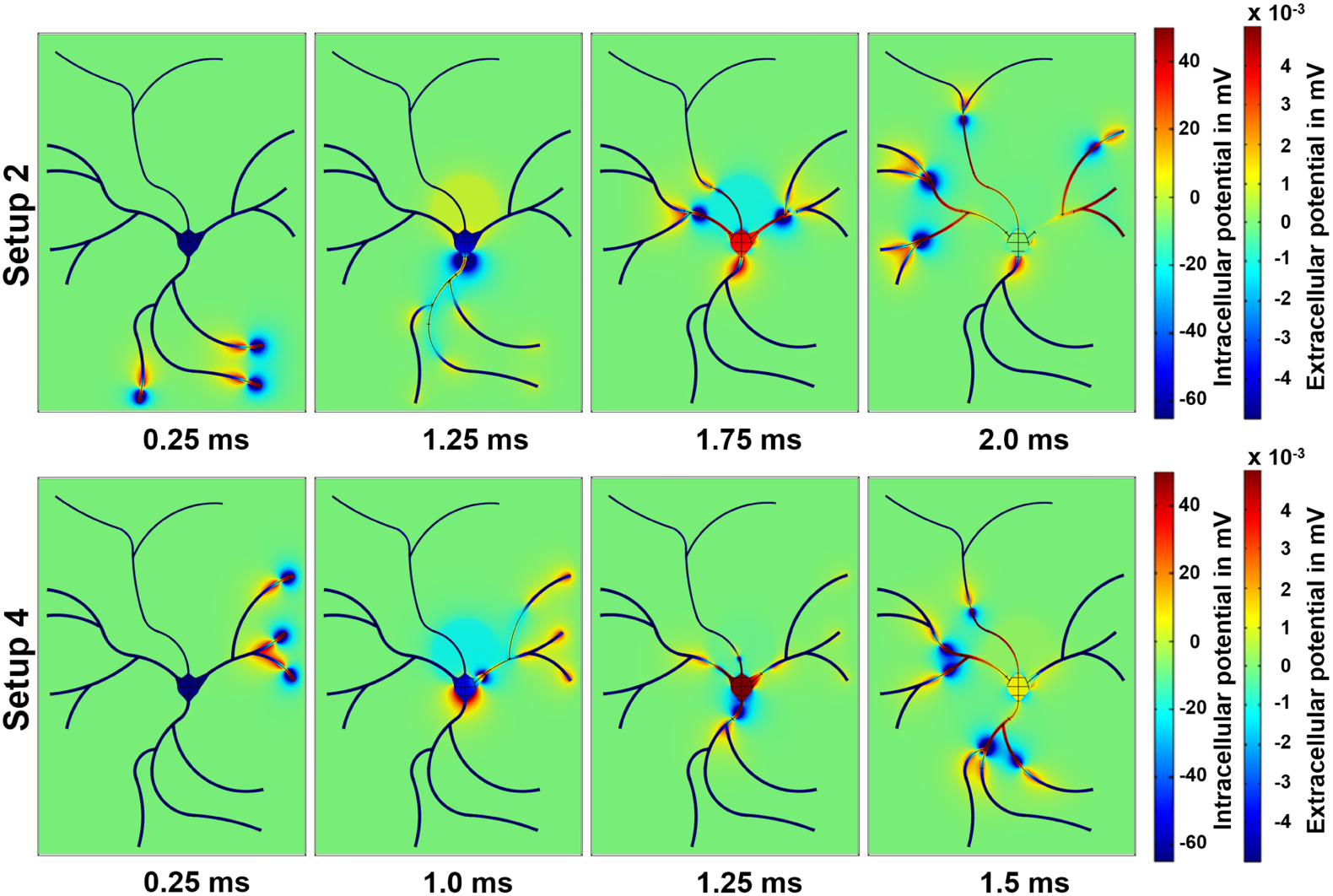
Intra- and extracellular potentials of scenarios 2 and 4 (top view). The origin of the stimulus in scenario 2 yields a positive potential at the electrode surface during the depolarization of soma and AIS, followed by a less pronounced negative potential in the subsequent repolarization phase. In contrast, in scenario 4, the potential distribution near the electrode consists mainly of an initial negative potential followed by a small positive potential distribution during the repolarization of soma and AIS.

Evoking AP-generation in the AIS via the basal dendrites, as in scenario 4, leads to a negative potential above the electrode while it causes a positive potential at the opposite side of the soma at the junction to the apical dendrite. Overall, the resulting extracellular potential distribution is similar to the results of models I-III without AP-stimulation via dendrites (see Fig. 6). In all these cases, AP-propagation through the soma yields a more pronounced *Na*^+^ current at the junction of soma to AIS and an increased *K*^+^ current near the junction to the apical dendrite (see Fig. A2). Therefore, the extracellular potential is negative near the AIS while it is positive at the junction between soma and apical dendrite.

In contrast, the distribution of the extracellular potential near the soma is reversed for scenario 2. The direction of the input potential that evokes AP-generation causes a more pronounced *Na*^+^ current into the soma at the junction to the apical dendrite and an increased *K*^+^ current near the axon hillock. Due to the changed distribution of the extracellular potential, this yields a substantially different electrode signal for identical recording positions.

In consequence, the location of the AP-evoking input is a vital factor that influences the resulting shape and amplitude of a derived MEA-electrode signal. Hence, this indicates that the structure of the neuronal network surrounding the measured cell may noticeably affect the result of the extracellular recording.

## IV. Discussion and Conclusion

Measuring the electrical activity of neurons using MEA-electrodes can yield a variety of different signal shapes, which can significantly hamper subsequent data analysis. In this context, the goal of this model-based study was identifying and analyzing the responsible factors.

Based on several FEM-models, multiple parameters with different influences on the resulting electrode signal were identified. Using model I, it was shown that parameters describing the electrical coupling between neurons and electrode primarily define the signal amplitude. The essential parameter was found to be the cell-coverage of the electrode, which can significantly in/decrease the signal amplitude by more than one order of magnitude. In addition, cell-coverage also influences the impact of other parameters, so that, e.g., the distance between neuron and electrode becomes a critical factor in case of sufficiently dense cell-coverage.

These findings also highlight the importance of an explicit simulation of the measuring electrode in order to accurately reproduce the effects of the neuron to electrode coupling. While methods like the point- and line-source approximation used in [9],[10] and [11] are suitable for an estimate of EAP at specific locations of extracellular space, such approximations do not account for interactions between the electrode and extracellular space. As shown using model I, there is indeed a very noticeable effect of a planar extracellular electrode on the extracellular potential in its vicinity. This is due to the fact that its presence alters the electric connection between the neuron, which works as a current source, and the electric reference potential as a current sink. In comparison, models using the point- and line-source approximation miss this effect as these entirely exclude the electrode from the model or only roughly describe it as an insulating surface.

Furthermore, models I, II, and III revealed a significant dependence of the derived signal shape on the electrode position. With model III, it was further shown that the path of incoming signals, which are used to evoke AP-generation, is an additional factor with substantial influence and noticeably changes the shape of derived electrode signals, even for identical positions. Our findings suggest that the shape of the extracellular recording not only contains information about the electrical activity of the measured neuron but also regarding the neuronal activity of neighboring cells. In consequence, information about the activity near the recording site is necessary to be able to analyze the geometric and morphologic situation at the respective electrode based on the measured electrical signal.

Together, all these results show that the relationship between neuron to electrode coupling and the derived electrode signal is very complex and dependent on a multitude of factors. Hence, the creation of a general overview of possible signal shapes is not possible. In addition, a detailed comparison between the results of different simulation models is not feasible, as any deviation in neuron geometry, morphology or even initial stimulus leads to different results. Similarly, comparing simulation to experimentally measured data can only yield meaningful insights, if factors like neuron geometry, electrode position and all the additional parameters that affect the neuron to electrode coupling are alike.

In this context, high-density microelectrode arrays offer the opportunity for further investigation. They allow for the derivation of neuronal signal shapes at multiple recording sites of a single neuron and therefore offer data about the path of AP-propagatiin as well as of local ion channel distribution.

Along with information about the neuron geometry, e.g. gathered through fluorescence staining, this data could be used to create a detailed mathematical model of a neuron that was previously measured using an HD-MEA setup. Once established this would allow for a detailed verification of the simulation model and furthermore may provide further findings concerning AP-generation, subsequent propagation and the formation of EAP.

## Appendix

All three gating variables are described by a differential equation that calculates their temporal evolution concerning empirically derived functions *α*_*k*_ and *β*_*k*_

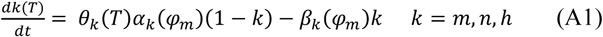

Both *α*_*k*_ and *β*_*k*_ depend on the transmembrane potential *φ*_*m*_ and are unique for each gating variable. The coefficient *θ*_*k*_ is used for temperature adaptation and calculated based on an empirical *Q*_10_ interval in the form of a scalar quantity as

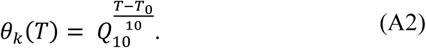

Besides, the ion-dependent permittivity *g*_*ion*_ changes with temperature

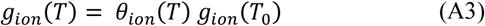

Based on the Nernst-equation, the reversal potential for each ion is affected

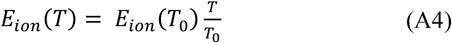

**Fig. A1.**
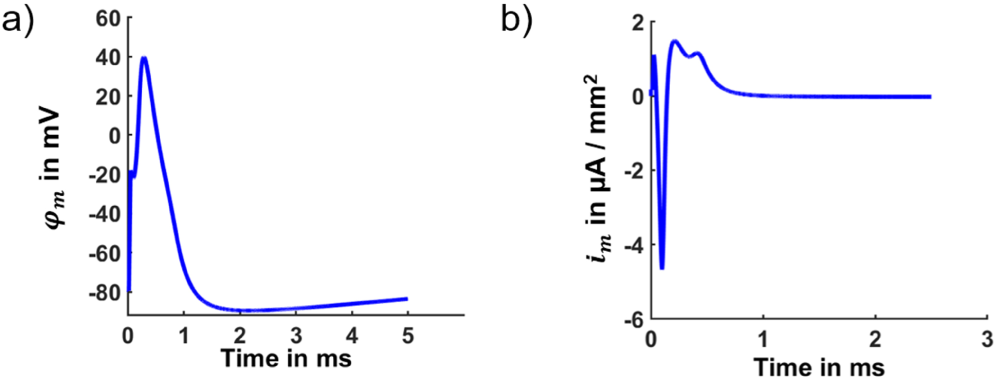
Transmembrane potential *φ*_*m*_ and respective transmembrane current density *i*_*m*_ of the stimulating signal used to evoke AP-generation and -propagation for model III.

**Fig. A2.**
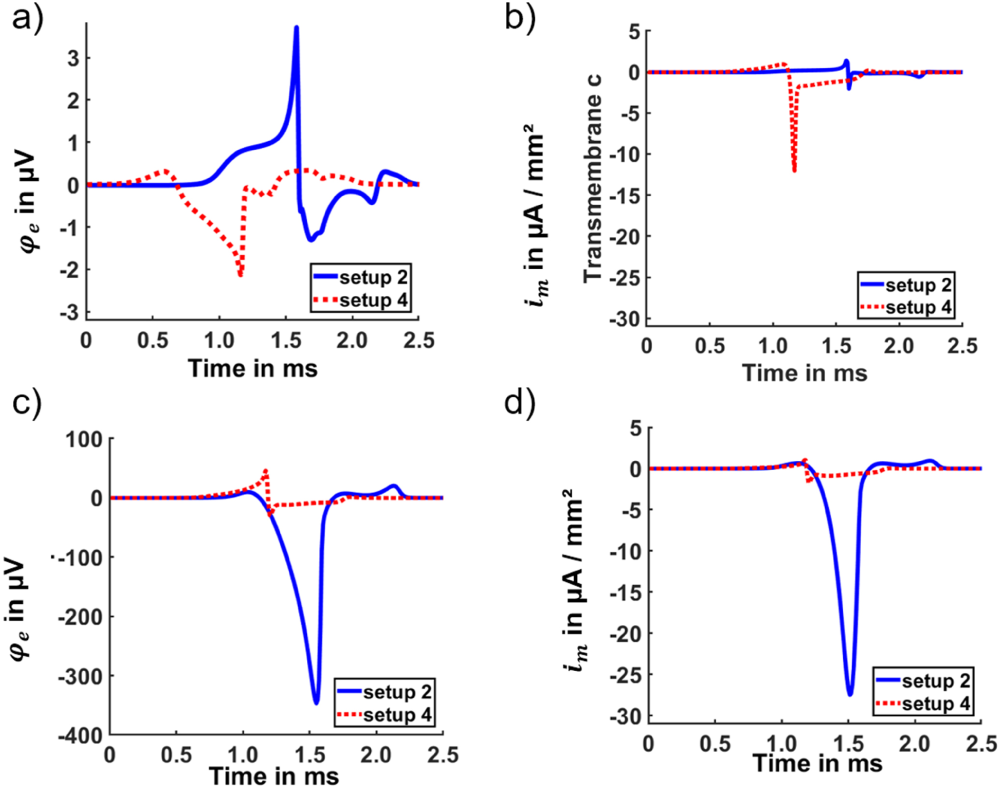
Resulting extracellular potential *φ*_*e*_ and respective transmembrane current density *i*_*m*_ at the axon hillock (a and b) and the junction of soma and dendrite (c and d) for stimulation scenarios 2 and 4 based on model III.

## Notes

This work was supported in part by the Bayrisches Staatsministerium für Bildung, Kultur, Wissenschaft und Kunst in the frame of ZEWIS and by Deutsche Forschungsgemeinschaft (DFG) under Grant AP 259/1-1.

### Competing Interest Statement

The authors have declared no competing interest.

